# Morning Engagement of Hepatic Insulin Receptors Improves Afternoon Hepatic Glucose Disposal and Storage

**DOI:** 10.1101/2024.09.25.614969

**Authors:** Hannah L. Waterman, Mary Courtney Moore, Marta S. Smith, Ben Farmer, Kalisha Yankey, Melanie Scott, Dale S. Edgerton, Alan D. Cherrington

## Abstract

Glucose tolerance improves significantly upon consuming a second, identical meal later in the day (second meal phenomenon). We previously established that morning hyperinsulinemia primes the liver for increased afternoon hepatic glucose uptake (HGU). Although the route of insulin delivery is an important determinant of the mechanisms by which insulin regulates liver glucose metabolism (direct hepatic vs indirect insulin action), it is not known if insulin’s delivery route affects the second meal response. To determine whether morning peripheral insulin delivery (as occurs clinically (subcutaneous)) can enhance afternoon HGU, conscious dogs were treated in the morning with insulin delivered via the portal vein, or peripherally (leg vein), while glucose was infused to maintain euglycemia. Consequently, arterial insulin levels increased similarly in both groups, but relative hepatic insulin deficiency occurred when insulin was delivered peripherally. In the afternoon, all animals were challenged with the same hyperinsulinemic-hyperglycemic clamp to simulate identical postprandial-like conditions. The substantial enhancement of HGU in the afternoon caused by morning portal vein insulin delivery was lost when insulin was delivered peripherally. This indicates that morning insulin does not cause the second meal phenomenon via its indirect actions on the liver, but rather through direct activation of hepatic insulin signaling.

**Article Highlights:**

- Morning insulin delivery primes the liver for increased hepatic glucose uptake (HGU) later in the day, but the mechanism (direct hepatic and/or indirect insulin action) remains unclear.
- This study compared insulin infusion via physiologic (hepatic portal vein) and clinical (peripheral) routes to assess their impact on afternoon hepatic glucose disposal.
- Morning peripheral insulin delivery failed to induce a significant enhancing effect on afternoon HGU and glycogen storage, unlike morning hepatic portal vein insulin delivery, which did.
- These findings highlight the importance of achieving appropriate hepatic insulin exposure in the morning to effectively prime the liver for efficient glucose disposal.

## Introduction

The liver plays a critical role in disposing of oral glucose by both suppressing glucose production and stimulating uptake (1; 2). This metabolic switch from fasting to feeding accounts for substantial meal-time glucose disposal in healthy individuals (3; 4). Given that the liver operates in storage mode two-thirds of the day and hepatic glucose uptake (HGU) and glycogen storage are impaired in people with diabetes, enhancing HGU could be a significant target for improving glycemic control and holds considerable therapeutic potential (5).

There is strong evidence that regularly consuming breakfast reduces the risk of obesity, cardiovascular disease, and type 2 diabetes (6). Yet, approximately 25% of individuals in the United States do not eat breakfast, and two-thirds frequently skip meals (7). The Staub-Traugott effect, or the second meal phenomenon, shows substantial improvement in glucose tolerance in response to a second identical meal (8–10). This occurs in both healthy individuals and those with diabetes, although the latter appears to have an abnormal response (11–13). Despite its identification over 100 years ago, the underlying mechanisms involved in this response remain unclear (14; 15).

We previously reported that morning (AM) hyperinsulinemia is essential for the second meal effect (16; 17). Remarkably, a physiologic rise in AM insulin doubled HGU and hepatic glycogen storage during an afternoon (PM) hyperinsulinemic-hyperglycemic clamp (16). To date, it is not known how insulin mediates this prolonged priming effect. However, it is well established that insulin acutely regulates hepatic glucose metabolism through both direct mechanisms (after binding to its hepatic receptors) and indirect mechanisms, acting at the brain (increasing neural signaling), adipose tissue (reducing circulating free fatty acids), and the pancreas (suppressing glucagon secretion) (18–20). Peripheral insulin delivery eliminates the physiological 3:1 insulin gradient between the hepatic portal vein and arterial circulation (generated by pancreatic insulin secretion into the portal vein) (21–25). Thus, when insulin is delivered subcutaneously, as is typical in individuals with diabetes, insulin’s direct effects on the liver are minimized while its indirect effects are amplified (21; 26). If the enhancement in PM HGU caused by hyperinsulinemia in the morning depends on insulin’s direct effects (favored by portal vein insulin delivery), then the disrupted distribution of insulin to the liver caused by AM peripheral delivery could impair the improved PM response, emphasizing the necessity of restoring the normal physiologic insulin gradient through therapeutic approaches such as delivering hepatopreferential or oral insulin analogs (22). Ultimately, this led us to investigate whether infusing insulin via its clinical route (i.e. peripheral delivery) in the morning would impact the liver’s ability to extract and store glucose later in the day.

## Research Design and Methods

### Animal care and surgical procedures

The study was conducted on 20 adult mongrel dogs (both male and female obtained from a USDA-licensed vendor, split evenly in each group) with an average weight of 25.1±0.7 kg. No significant sex differences were observed for any parameter measured. The dogs were cared for per the American Association for the Accreditation of Laboratory Animal Care standards. The experimental protocol was approved by the Vanderbilt Institutional Animal Care and Use Committee. Two weeks before each experiment, a laparotomy was performed under general anesthesia to position blood flow probes around the hepatic artery and hepatic portal vein in the dog. Sampling catheters were implanted in the hepatic vein, hepatic portal vein, and femoral artery, and infusion catheters were placed into vessels feeding into the hepatic portal vein (the splenic and jejunal veins), and inferior vena cava. These catheters were secured subcutaneously until the study day. Dogs were fed a diet of chow and meat with 46% carbohydrate, 34% protein, 14.5% fat, and 5.5% fiber, and were fasted for 18 hours before the experiment. Only healthy dogs were included in the study, based on meal consumption (>75% of their last meal), leukocyte count (<18,000/mm³), and hematocrit levels (>34%). Blood withdrawal was limited to 20% of total blood volume.

### Experimental Design

#### Morning (AM) Clamp Period (0-240 min)

The experimental protocol comprised two glucose clamping periods representative of breakfast and lunch (AM and PM), with an interval of rest between them **(Fig. 1)**. Blood was collected every 15-30 minutes from the femoral artery, hepatic portal vein, and hepatic vein for hormone and substrate analysis, and arterial plasma glucose was monitored every 5 minutes.

**Figure 1:**
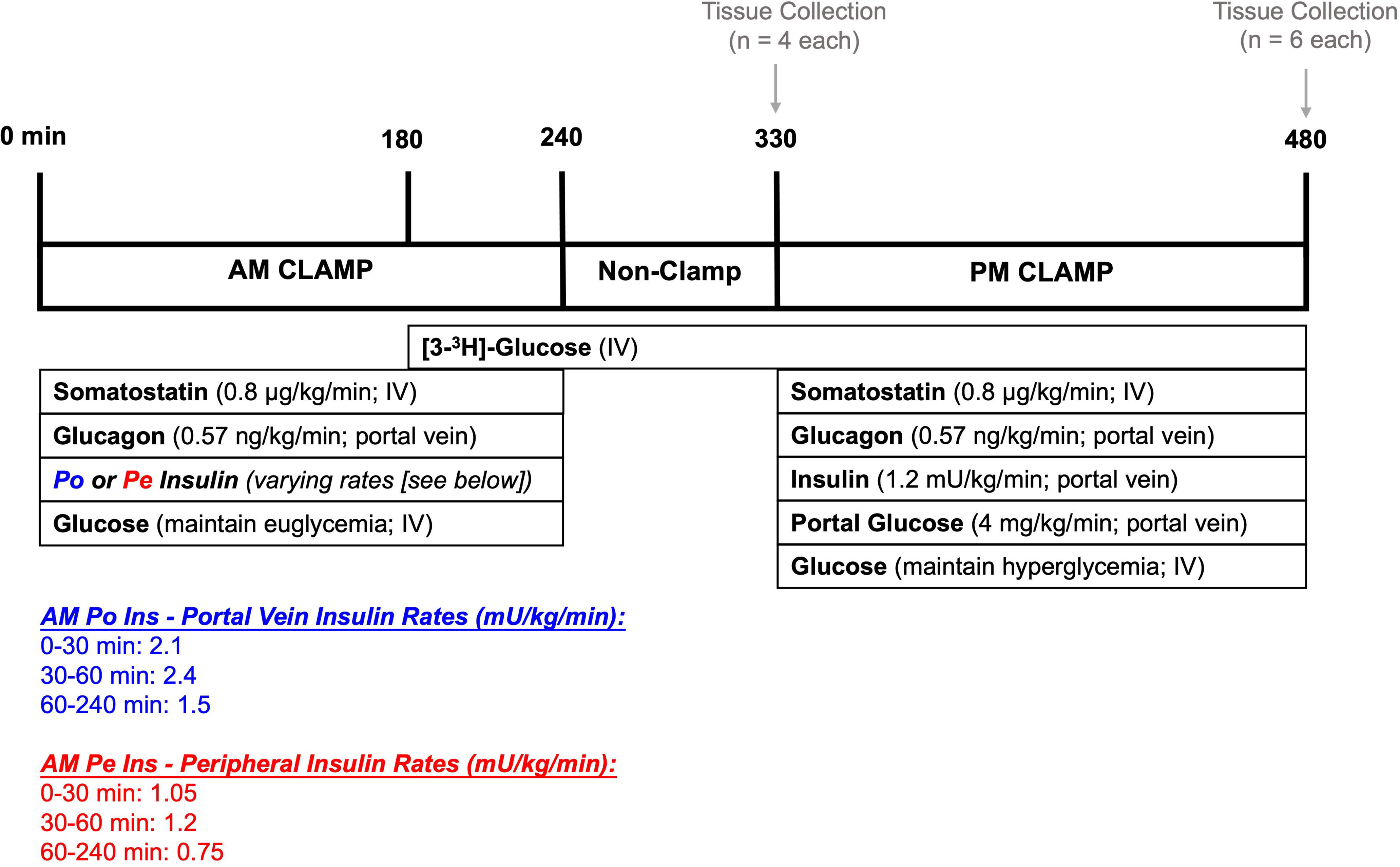
Experimental protocol. Canines underwent a 4h euglycemic clamp in the morning (0 to 240 min) with either hepatic portal vein insulin infusion (Group 1: AM Po Ins) or leg vein peripheral insulin infusion (Group 2: AM Pe Ins) at rates designed to match circulating arterial insulin levels and create a differential effect at the liver. Near the end of the AM clamp, tracer infusion began to allow enough time for equilibration before the start of the PM clamp (180 min). Following the AM clamp, there was a 1.5h rest period (240-330 min) where all infusions were halted. At the end of this rest period, dogs from each group were euthanized and hepatic tissue was collected for molecular analysis (*n*=4/group). The remainder of the dogs (*n*=6/group) underwent a 2.5h PM hyperinsulinemic-hyperglycemic clamp with portal glucose delivery (330 to 480 min). Tissue collection occurred at the end of the PM clamp for these dogs. Details can be found in the methods section. IV; intravenous infusion.

Having previously established that insulin (not glucose) is the critical factor during the morning (16), we performed a hyperinsulinemic-euglycemic morning clamp. At the onset of this period, somatostatin (Bachem, Torrance, CA) was infused via the inferior vena cava (0.8 µg/kg/min) to suppress endogenous pancreatic insulin and glucagon secretion. Glucagon (GlucaGen, Boehringer Ingelheim, Ridgefield, CT) was replaced intraportally (0.57 ng/kg/min) to maintain its level at basal. One group (AM Po Ins) received AM insulin (Novolin R; Novo Nordisk, Basværd, Denmark) infusion directly into the hepatic portal vein to replicate the physiologic pattern of insulin secretion that was previously observed in response to an AM duodenal glucose infusion (2.1 mU/kg/min [0-30 min], 2.4 mU/kg/min [30-60 min], and 1.5 mU/kg/min [60-240 min]) (16). The other group (AM Pe Ins) received one-half the amount of insulin that the AM Po Ins group received, but it was delivered into a leg vein rather than intraportally (1.05 mU/kg/min [0-30 min], 1.2 mU/kg/min [30-60 min], and 0.75 mU/kg/min [60-240 min]). These rates were chosen to account for ∼50% hepatic insulin extraction so that, by design, arterial insulin levels would be matched between groups, while hepatic insulin levels would be reduced in the AM Pe Ins group, as occurs when insulin is delivered via the peripheral route (20; 22). Glucose was infused into a leg vein and adjusted as needed to clamp arterial blood glucose at euglycemia (plasma glucose at ∼100 mg/dL). To allow for the calculation of HGU during the afternoon period, peripheral [3-^3^H] glucose (Revvitty, Waltham, MA) infusion began at 180 min in both groups (38 µCi prime and 0.38 μCi/min continuous rate). At the end of the AM clamp (240 min), all infusions besides the glucose tracer were halted.

#### Non-clamp Period (240 – 330 min)

Blood samples were collected from the arterial, portal vein, and hepatic vein sampling catheters at 300, 315, and 330 min to allow assessment of glucose kinetics before the onset of the afternoon (PM) clamp **(Fig. 1)**. To determine the liver’s molecular status at the start of the PM period, a subset of 4 dogs from each group were anesthetized at 330 min, hepatic tissue was quickly collected and flash-frozen in liquid nitrogen to preserve cellular conditions, then the animals were euthanized. These samples were stored at −80°C. The remainder of the dogs (n=6/group) underwent the PM clamp.

#### Afternoon (PM) Clamp Period (330 – 480 min)

The AM Po Ins and AM Pe Ins groups both underwent the same 2.5h PM hyperinsulinemic-hyperglycemic clamp **(Fig. 1)**. Somatostatin was infused as described above, while basal glucagon (0.57 ng/kg/min) and 4x basal insulin (1.2 mU/kg/min) were infused intraportally. Additionally, glucose was infused into the hepatic portal vein to activate the portal glucose feeding signal (4 mg/kg/min) (5). A primed, continuous infusion of glucose was delivered into a leg vein and adjusted as necessary to maintain 2-fold basal hyperglycemia throughout the PM clamp (plasma glucose at ∼200 mg/dL). These clamp conditions resemble the physiologic environment observed after the consumption of a carbohydrate-rich meal, at a steady state (5). Upon collecting the final blood sample (480 min), the dogs were anesthetized, and hepatic tissue was collected and stored as described above.

### Analyses

#### Biochemical and Molecular Methods

Whole blood samples were analyzed to determine hormone and substrate balance across the liver using standard methods (27). During the experimental period, plasma glucose levels were immediately measured using an Analox GM9 glucose analyzer. Insulin (#PI-12K, MilliporeSigma, Burlington, MA), glucagon (#GL-32K, MilliporeSigma), and cortisol (VUMC Analytical Services in-house primary antibody with I^125^ cortisol from MP Biomedicals, Santa Ana, CA) were measured by radioimmunoassay (27). Metabolites involved in key metabolic processes, including lactate, glycerol, alanine, and non-esterified fatty acids (NEFA) were assessed using enzymatic spectrophotometric methods (27). To determine the amount of [3-^3^H]-glucose in each sample, plasma was deproteinized and quantified using liquid scintillation counting to measure tracer-specific activity (28; 29). Hepatic glycogen was quantitatively assessed using the Keppler and Decker amyloglucosidase method (30).

All molecular analysis was performed on terminal hepatic tissue samples, including qPCR analysis (glucokinase [GK] mRNA), western blotting (Akt, glycogen synthase [GS], glycogen phosphorylase [GP], and GK proteins), and enzymatic activity assays using radioisotope and colorimetric methods (GS, GP, and GK) (27). Quantities of transcripts were normalized to that of the housekeeping gene GAPDH, and proteins were normalized to either total protein or cyclophilin B, as appropriate. The molecular methods were optimized to assess canine-specific proteins using validated primers and antibodies (31). Baseline liver samples from five overnight-fasted dogs were included for reference.

#### Calculations

Hepatic glucose load was determined using the following equation: [HGLin] = [BFa x Ga + BFp x Gp], where G represents the blood glucose concentration, BF represents measured blood flow, and A, P, and H represent the hepatic artery, hepatic portal vein, and hepatic vein. Hepatic sinusoidal plasma insulin and glucagon concentrations were calculated based on the percent contribution of the hormones in the hepatic artery and portal vein (27). Unidirectional hepatic glucose uptake (HGU), non-HGU, and direct glycogen synthesis (glycogen synthesized from glucose) were calculated using [3-^3^H]-glucose as previously described, where non-HGU is the difference between net-HGU and the glucose infusion rate, adjusted for glucose mass changes (27). Net hepatic glucose balance (NHGB), net hepatic carbon retention (NHCR), indirect glycogen synthesis (glycogen synthesized from gluconeogenic precursors), and gluconeogenic flux were determined as previously outlined (27; 32; 33). Hepatic fractional extraction of insulin was calculated by dividing net hepatic insulin balance by the inflowing arterial and portal vein plasma insulin concentrations. Insulin-independent glucose uptake was estimated to be 67% (approximately 1 mg/kg/min) of net hepatic glucose output in the basal state (34).

#### Statistics

All data are presented as mean ± SEM. Statistical comparisons between groups and over time were performed using a two-way ANOVA with repeated measures design with Student-Newman-Keuls multiple comparisons post hoc analysis. For respective areas under the curve, a one-way ANOVA with Tukey’s post hoc analysis was used. P<0.05 was considered statistically significant. GraphPad Prism software was utilized for all statistical analysis.

#### Data and Resource Availability

Values for all data points in graphs are reported in the Supporting Data Values file. Additionally, these data and supplemental materials are publicly available via the following DOI for Figshare: doi.org/10.6084/m9.figshare.26800804. All other data generated and analyzed during the current study are available from the corresponding author upon reasonable request.

## Results

### AM Clamp Data

Euglycemia was successfully maintained throughout the AM clamp without significant group differences **(Fig. 2A).** As intended, arterial plasma insulin levels increased similarly in both groups (53 ± 3 and 47 ± 2 µU/mL in the AM Po Ins and AM Pe Ins groups, respectively; **Fig. 2B-C**; **Table 1**). Portal vein insulin infusion raised hepatic sinusoidal insulin levels significantly (to 137 ± 11 µU/mL in AM Po Ins), which was 2.6-fold greater than the arterial insulin levels **(Fig. 2B-E**, **Table 1).** In contrast, with peripheral vein insulin infusion, this physiologic increase in insulin at the liver did not occur **(Fig. 2B-E**, **Table 1).** The glucose infusion rate (GIR) to maintain euglycemia was slightly but not significantly higher in the AM Po Ins group **(Fig. 2F, 2H).** All animals were in a state of hepatic glucose production by the onset of the PM clamp and glucose levels, circulating NEFAs, and metabolite levels (amino acids, glycerol, and lactate) had returned to baseline.

**Figure 2:**
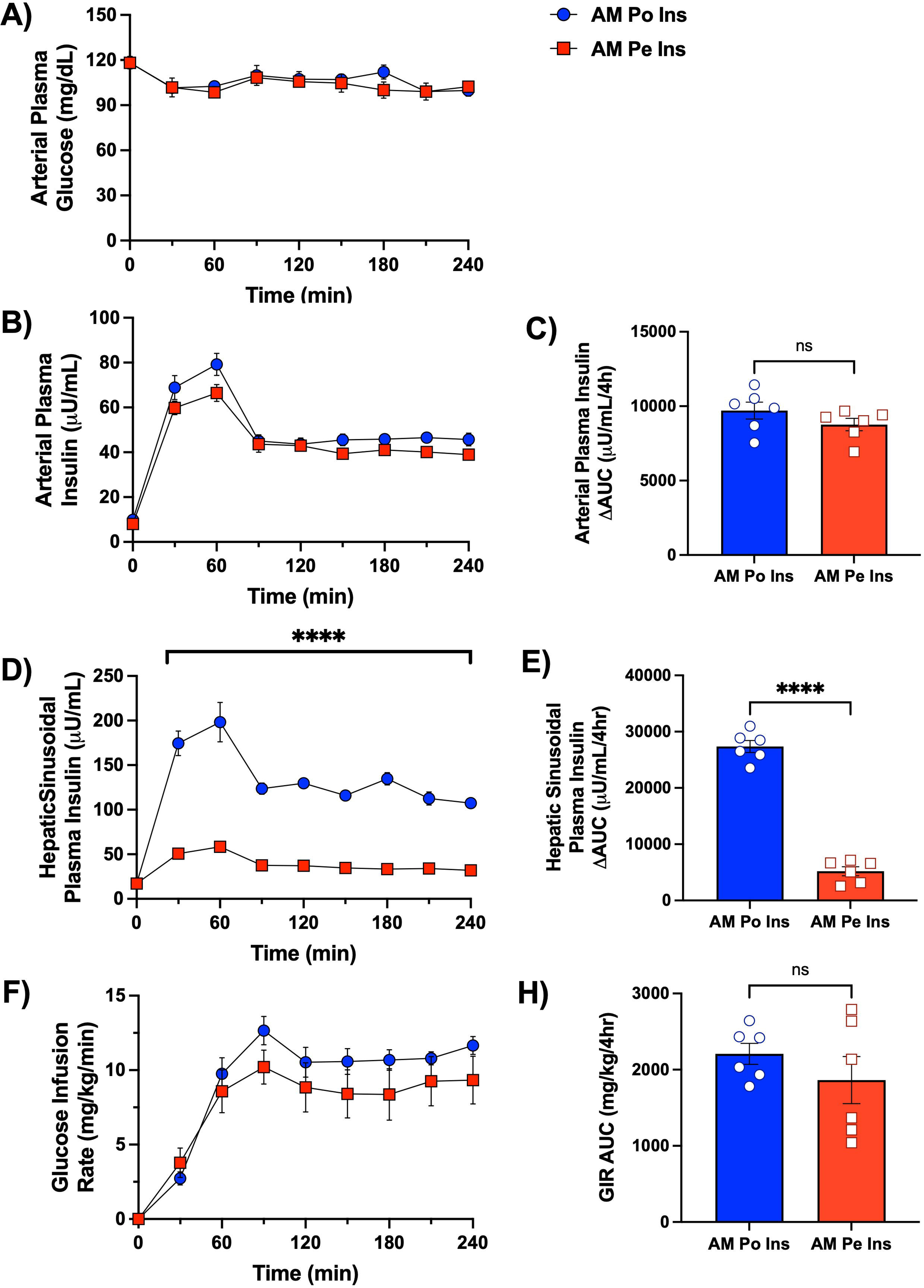
Morning (AM) clamp glucose and insulin flux data. Arterial plasma glucose (A) and insulin (B), hepatic sinusoidal plasma insulin (D), and the amount of exogenous glucose required to maintain euglycemia (F) are shown for the AM Po Ins and AM Pe Ins group; *n*=6/group. Bar graphs indicate the area under the curve (AUC) for arterial insulin concentrations (C), hepatic sinusoidal insulin concentrations (E), and glucose infusion rates (H) during the 4h morning period. Data are expressed as mean ± SEM. ****P<0.0001 between groups.

**Table 1:**
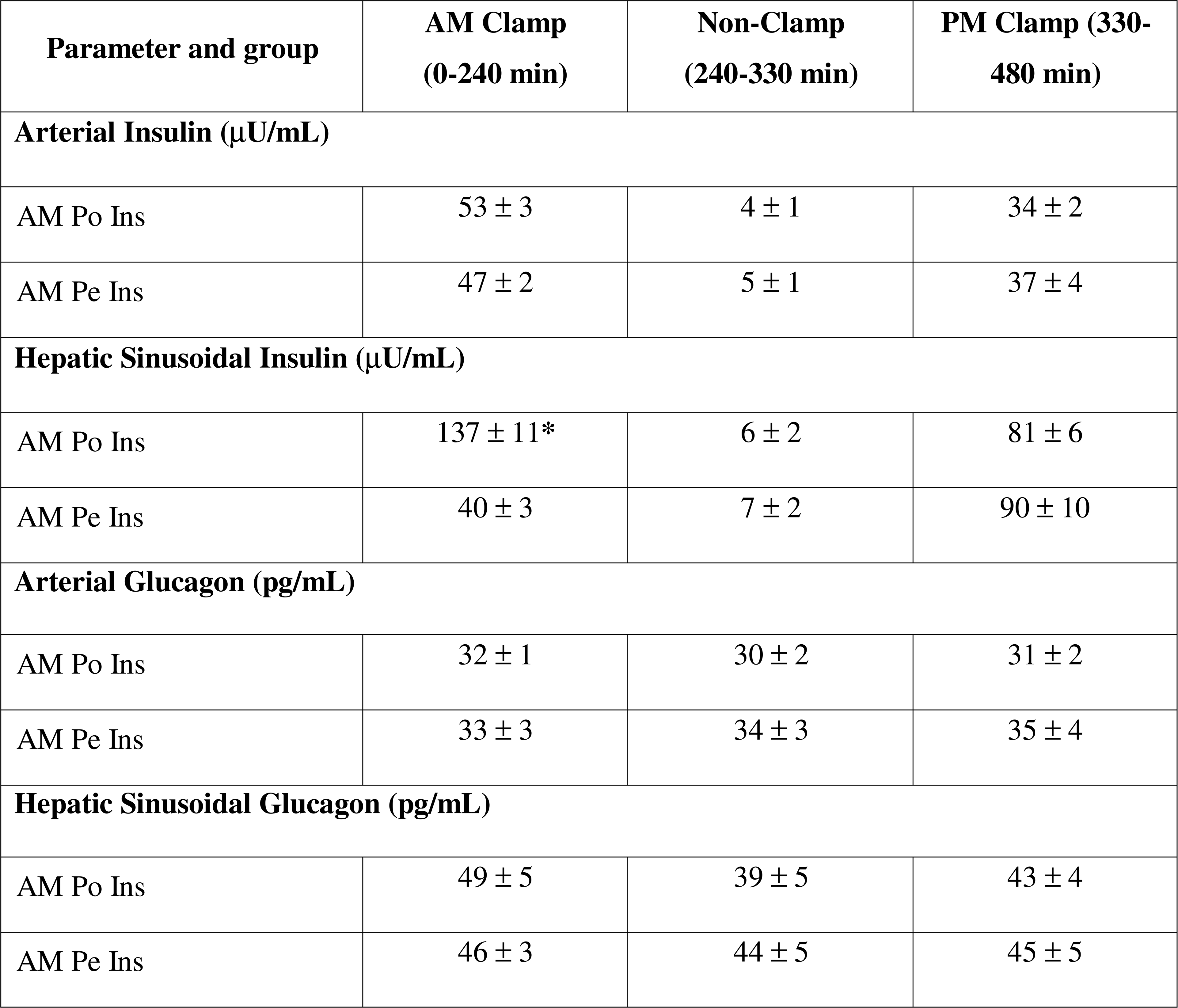
Average plasma hormone concentrations throughout the experimental protocol. Insulin samples were analyzed every 30 min and glucagon samples were analyzed every hour during the experimental protocol. Each value represents the mean of all samples collected within each clamping period. Data are represented as mean ± SEM, n=6 per group, *P<0.05.

| Parameter and group | AM Clamp<br>(0-240 min) | Non-Clamp<br>(240-330 min) | PM Clamp (330-480 min) |
| --- | --- | --- | --- |
| **Arterial Insulin ($\mu\text{U/mL}$)** |  |  |  |
| AM Po Ins | $53 \pm 3$ | $4 \pm 1$ | $34 \pm 2$ |
| AM Pe Ins | $47 \pm 2$ | $5 \pm 1$ | $37 \pm 4$ |
| **Hepatic Sinusoidal Insulin ($\mu\text{U/mL}$)** |  |  |  |
| AM Po Ins | $137 \pm 11^*$ | $6 \pm 2$ | $81 \pm 6$ |
| AM Pe Ins | $40 \pm 3$ | $7 \pm 2$ | $90 \pm 10$ |
| **Arterial Glucagon (pg/mL)** |  |  |  |
| AM Po Ins | $32 \pm 1$ | $30 \pm 2$ | $31 \pm 2$ |
| AM Pe Ins | $33 \pm 3$ | $34 \pm 3$ | $35 \pm 4$ |
| **Hepatic Sinusoidal Glucagon (pg/mL)** |  |  |  |
| AM Po Ins | $49 \pm 5$ | $39 \pm 5$ | $43 \pm 4$ |
| AM Pe Ins | $46 \pm 3$ | $44 \pm 5$ | $45 \pm 5$ |

### PM Clamp Data

Postprandial conditions were achieved during the PM clamp by doubling blood glucose levels **(Fig. 3A, 3B, 3D),** establishing a negative arterial-to-portal vein glucose gradient **(Fig. 3C),** and quadrupling arterial and hepatic insulin levels **(Fig. 3E, 3F)** via intraportal glucose and insulin infusions in both groups (5). Plasma glucagon concentrations were maintained at basal levels **(Table 1)**, and cortisol was low in both groups, indicating minimal stress. No group differences in key liver glucose metabolism determinants were observed.

**Figure 3:**
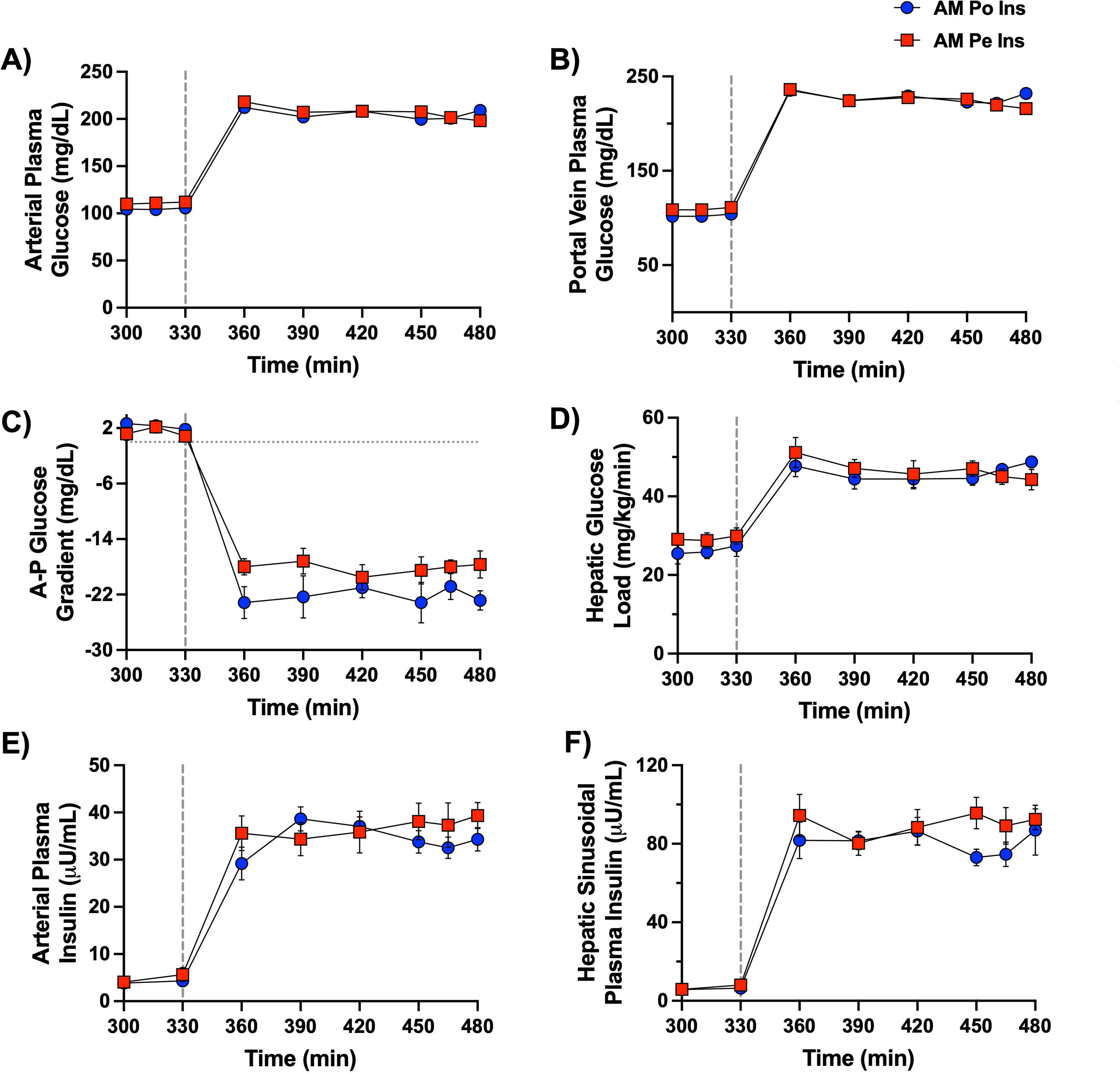
Afternoon (PM) clamp glucose and hormone data. A vertical line at 330 min separates the resting period from the onset of the PM clamp. Arterial plasma glucose (A), portal vein plasma glucose (B), the difference between the artery and the portal vein plasma glucose levels (C), hepatic glucose load (D), arterial plasma insulin (E), and plasma insulin at the hepatic sinusoids (F) are shown for the 2.5h hyperinsulinemic-hyperglycemic PM clamp period in both the AM Po Ins and AM Pe Ins group; *n*=6/group. There were no differences between the two groups for any of the parameters measured. Data are expressed as mean ± SEM.

Despite exposure to the same PM clamp conditions, the GIR was greater in the AM Po Ins group compared to the AM Pe Ins group (mean PM GIR of 13.8 ± 1.2 vs. 9.5 ± 1.0 mg/kg/min, respectively, P<0.05, **Fig. 4A, 4B**). This difference in PM GIR was not due to a significant increase in PM non-hepatic glucose uptake (AUC of 972 ± 130 vs. 825 ± 94 mg/kg/2.5h in AM Po Ins vs. AM Pe Ins, respectively, **Fig. 4C, 4D**). Rather, the difference was primarily due to a remarkable near 2-fold enhancement in PM HGU (mean 6.3 ± 0.5 in AM Po Ins vs. 3.5 ± 0.3 mg/kg/min in AM Pe Ins, **Fig. 4E, 4F**). Of the total amount of glucose disposed of in the AM Po Ins group during the PM clamp, glucose uptake was distributed similarly between liver and muscle (AUCs of 830 ± 79 and 760 ± 117 mg/kg/2.5h, respectively), with insulin-independent tissues disposing of 12% of total GIR (**Fig. 5)**. In contrast, in the AM Pe Ins group, total glucose disposal was reduced by 31% (AUC of 1238 ± 124 mg/kg/2.5h), HGU was down 50% (AUC of 413 ± 54 mg/kg/2.5h), MGU decreased 18% (AUC of 627 ± 90 mg/kg/2.5h), and insulin-independent tissue disposal remained similar compared to the AM Po Ins group (**Fig. 5).** Instead of the equal HGU to MGU percent distribution of PM glucose disposal that occurred in the AM Po Ins group, MGU was 1.5x that of HGU in the AM Pe Ins group.

**Figure 4:**
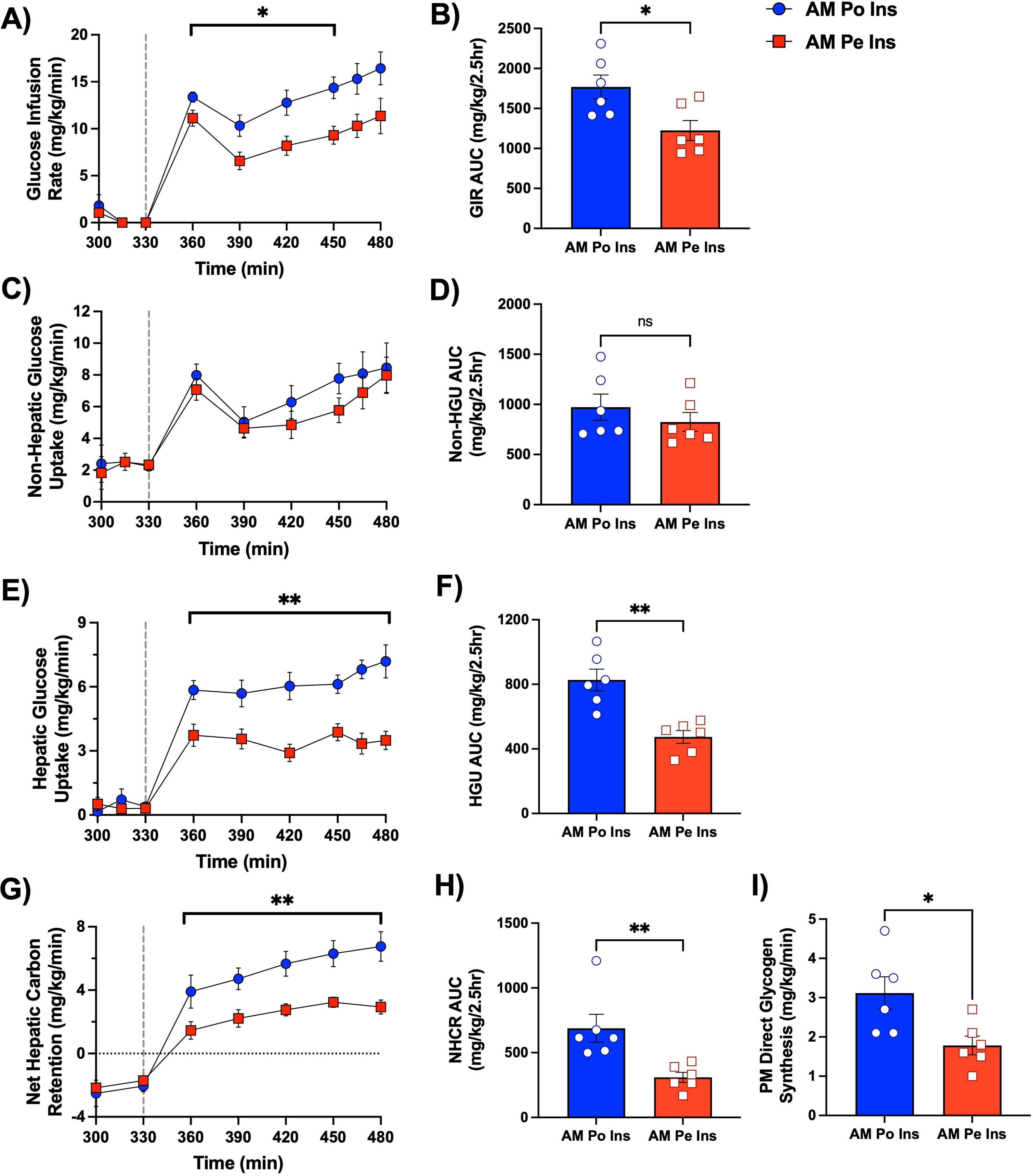
Glucose uptake and storage during the PM hyperinsulinemic-hyperglycemic clamp. A vertical line at 330 min separates the resting period from the onset of the PM clamp. Glucose infusion rate (A), non-hepatic glucose uptake (C), hepatic glucose uptake (E), and net hepatic carbon retention (G) are shown over time for the AM Po Ins and AM Pe Ins group, *n*=6/group. The respective AUCs for each of these measurements are shown for both groups (B, D, F, H). Tracer-determined direct glycogen synthesis (I) is shown for both groups, as well. *P<0.05, **P<0.01 between groups. Data are expressed as mean ± SEM.

**Figure 5:**
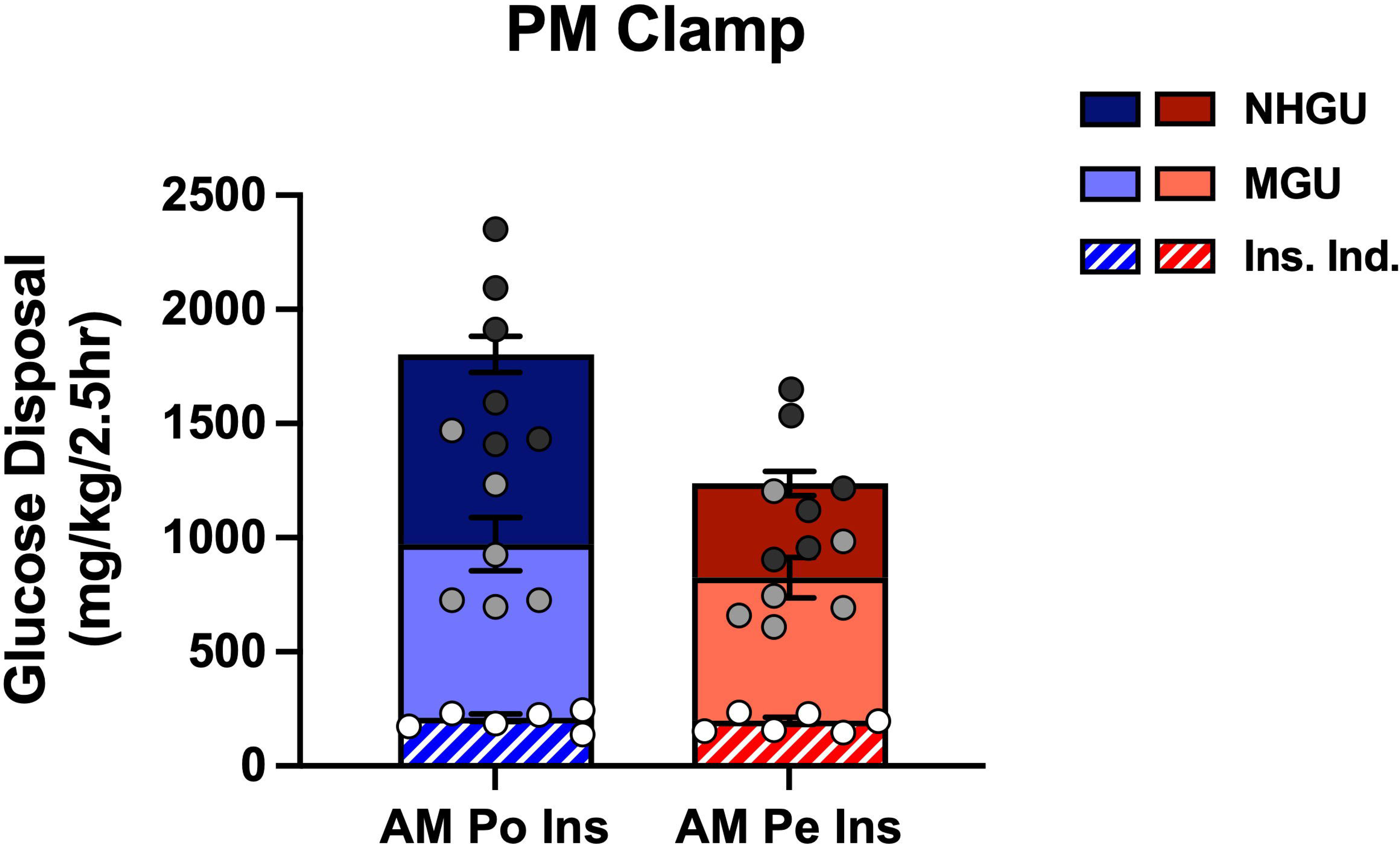
Distribution of glucose disposal during the PM clamp. Each bar graph represents the total amount of glucose infused in each group during the PM clamp. This is further divided into the mean amount of glucose being taken up by the liver (NHGU), skeletal muscle (MGU), and insulin-independent tissues (Ins. Ind.), such as the brain, smooth muscle, and red blood cells. Individual points represent Ins. Ind. glucose uptake (white dots), Ins. Ind. and MGU (non-HGU; light grey dots), and total glucose disposal (total glucose infused (Ins. Ind, MGU, and NHGU together); dark grey dots). Bar graph data are expressed as mean ± SEM.

During the PM clamp, net hepatic carbon retention (NHCR; this rate includes glucose, amino acids, glycerol, and lactate taken up by the liver) was markedly augmented in the AM Po Ins group (mean PM NHCR of 5.3 ± 0.5 mg/kg/min) in comparison to the AM Pe Ins group (mean PM NHCR of 2.5 ± 0.3 mg/kg/min) **(Fig. 4G, 4H).** PM Direct glycogen synthesis (the rate of radiolabeled plasma glucose incorporated into hepatic glycogen reserves) was significantly greater with morning portal vs. peripheral vein insulin infusion (**Fig. 4I)**. Throughout the PM clamp, a total of 26 ± 3 mg glycogen/g liver was synthesized by the direct pathway in the AM Po Ins group, 1.6-fold more than in the AM Pe Ins group (16 ± 1 mg glycogen/g liver, **Fig. 4I**). At the onset of the PM clamp, hepatic glycerol and NEFA uptake were quickly suppressed, and all animals switched to a state of net hepatic lactate output **(Fig. 6)**. Glycolytic flux was greater in the AM Po Ins vs. the AM Pe Ins group (mean glycolytic flux of 1.7 ± 0.1 vs. 1.2 ± 0.1 mg/kg/min, respectively, data not shown), supporting the observed increase in net hepatic lactate output in the AM Po Ins group.

**Figure 6:**
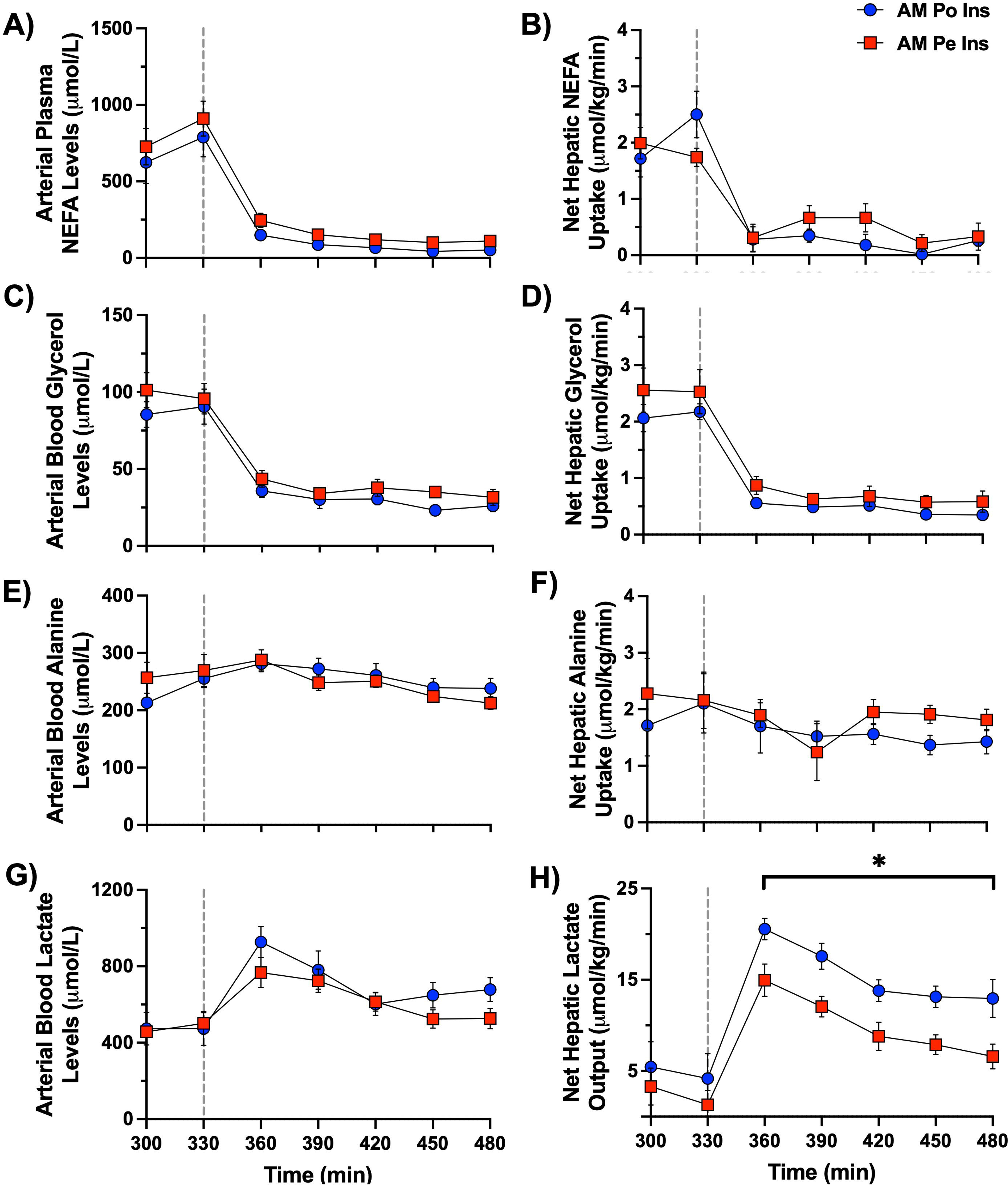
PM clamp fatty acid and metabolite flux data. A vertical line at 330 min separates the resting period from the onset of the PM clamp. Arterial plasma non-esterified fatty acids (A) and net hepatic uptake (B) are shown for the 2.5h PM clamp. Arterial blood glycerol (C), alanine (E), and lactate (G) levels are shown along with their respective net hepatic balance (D, F, H) for the AM Po Ins and AM Pe Ins group, *n*=6/group. *P<0.05 between groups. Data are expressed as mean ± SEM.

### Liver tissue analyses

When assessing Akt phosphorylation, there were no differences between basal conditions and either group prior to the PM clamp, while both groups were similarly elevated at the end of the PM clamp (**Fig. 7A**). This indicates that Akt signaling, activated by insulin in the morning, returned to baseline by the end of the 90 min rest period in both groups. On the other hand, morning portal vein insulin delivery had a pronounced effect on hepatic GK gene transcription, resulting in GK mRNA levels that were 3.5-fold higher in the AM Po Ins group vs. the AM Pe Ins group at the start of the PM clamp **(Fig. 7B)**. GK mRNA levels did not increase further post-PM clamp **(Fig. 7B)**. The changes observed with GK transcript levels correlated closely with GK protein in both groups, suggesting that increased GK during the PM clamp was concomitant with the increase in HGU observed in the AM Po Ins group **(Fig. 7C)**.

**Figure 7:**
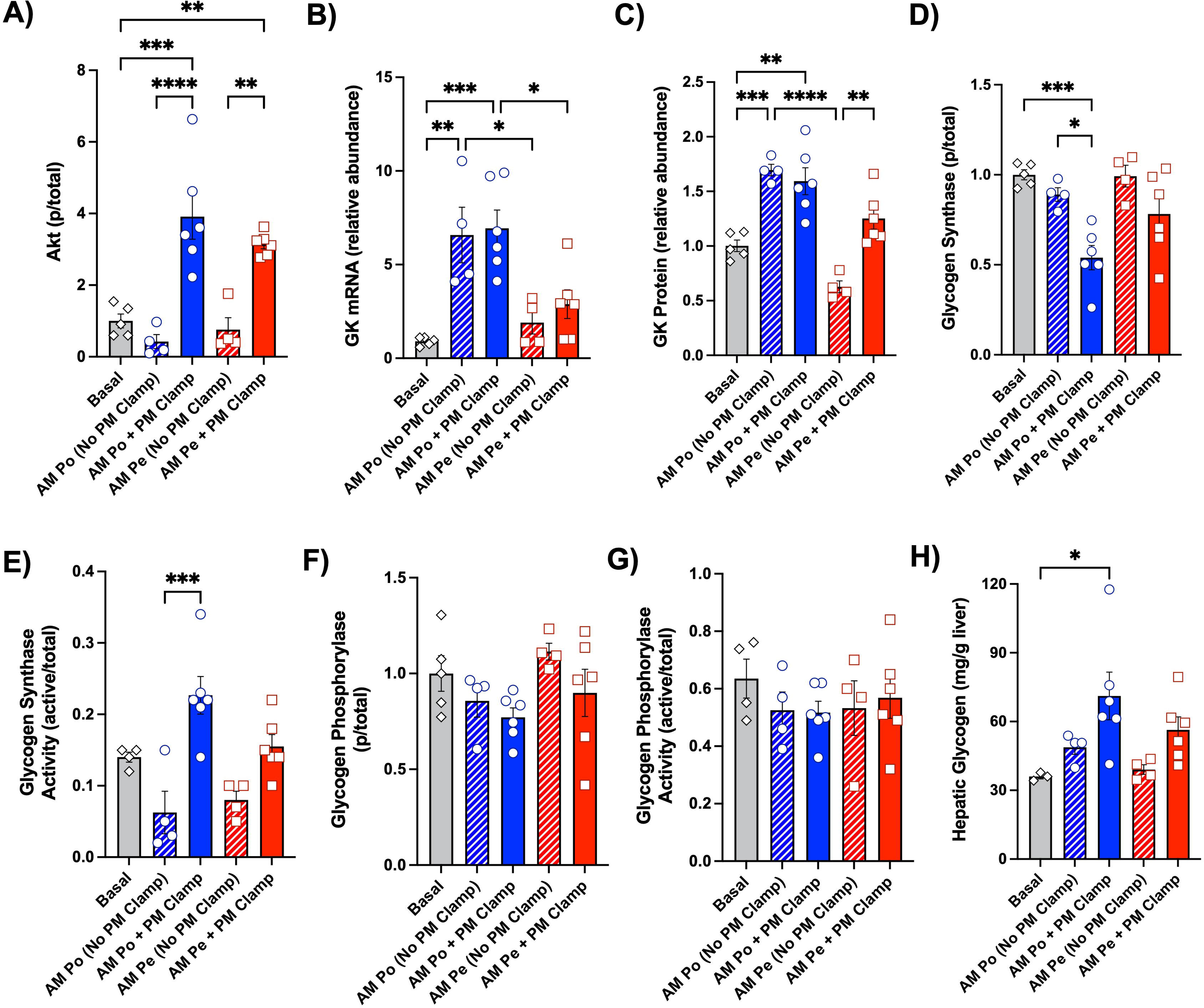
Hepatic tissue molecular analyses. Phosphorylated Akt protein (A), glucokinase (GK) mRNA (B), GK protein (C), phosphorylated glycogen synthase (GS) protein (D), GS activity (E), phosphorylated glycogen phosphorylase (GP) protein (F), GP activity (G), and terminal hepatic glycogen (H) are shown for basal samples (*n*=3-5), the AM Po Ins group (*n*=6), and the AM Pe Ins group (*n*=6). Data are expressed as mean ± SEM. *P<0.05, **P<0.01, ***P<0.001, ****P<0.0001 between relevant groups. All other comparisons not denoted with a P-value are not significant (ns)

Regarding glycogen synthase (GS), no differences in phosphorylation or activation were detected between basal tissue and liver samples collected prior to the PM clamp for either group **(Fig. 7D, 7E)**. The most substantial effect on GS was observed in the AM Po Ins group, where there was significantly greater dephosphorylation following the PM clamp, resulting in increased GS activity. There was a tendency for greater GS dephosphorylation and activation in the AM Pe Ins group at the end of the PM clamp, although statistical significance was not reached (P<0.1, **Fig. 7D, 7E)**. As for glycogen phosphorylase (GP) phosphorylation and activation, there were no large differences between any of the groups **(Fig. 7F, 7G)**. The molecular changes observed with GK and GS correspond nicely with the rate for tracer-direct glycogen synthesis in each group and measured terminal hepatic glycogen levels **(Fig. 4I, 7H**). Thus, it appears that the greater increase in AM direct hepatic insulin action better primed the liver for greater activation of GK and GS in the PM, which through a coordinated effort, assisted in the enhancement of PM hepatic glucose disposal and hepatic glycogen synthesis.

## Discussion

Although the second meal phenomenon was first observed early into the 20^th^ century, there remains only rudimentary knowledge regarding the physiologic and cellular mechanisms that explain this effect (14; 15). We recently demonstrated that improved glucose tolerance during the second meal is linked to AM hyperinsulinemia prompted by a morning glucose load (16). While insulin regulates the liver directly, the hormone also affects other insulin-sensitive organs (e.g., brain, fat, and pancreas) that indirectly control hepatic glucose metabolism (19; 35-38). Peripheral insulin delivery eliminates the normal, physiological insulin gradient between the liver and the rest of the body (21–25). Consequently, the relative hepatic insulin deficiency that inevitably results from peripheral insulin delivery shifts the balance of control of liver glucose metabolism away from direct mechanisms, towards indirect regulators of the liver (39). Given that the route of insulin delivery is therefore a determinant of the mechanisms by which insulin acts on the liver, we aimed to establish whether the distribution of insulin in the circulation affects how the liver is primed in the morning for enhanced HGU later in the day.

We found that afternoon HGU and glycogen storage were only enhanced when there was a sufficient increase in insulin at the liver (portal vein delivery) in the morning. Remarkably, HGU was nearly 2-fold greater, with most of the glucose being stored as glycogen, without a significant difference in PM muscle glucose uptake. Thus, the direct action of insulin (binding to hepatocytes) in the morning resulted in what was likely a series of cellular changes that prepared the liver to better respond to a second glucose load by changing the liver’s molecular setpoint. Together, these data demonstrate that without a concomitant 2.5-fold greater increase in insulin at the liver, a morning increase in arterial insulin (as occurs with subcutaneous insulin injection), was unable to improve hepatic glucose handling later in the day.

There are several mechanisms by which insulin’s indirect effects in the morning could have mediated its PM priming effect. First, when considering its effects on the brain, it has been demonstrated in rodents and canines that a physiologic increase in brain insulin action activates hypothalamic Akt, alters STAT3 signaling in the liver, significantly increases GK and glycogen synthase kinase 3 beta (GSK3β) gene expression, and suppresses the transcription of hepatic gluconeogenic genes (37; 38; 40-43). In the dog, these changes led to an increase in NHGU that only manifested after prolonged (>3h) exposure of the brain to insulin (38). Thus, it is reasonable to postulate that 4h of AM brain insulin action could prime the liver for enhanced PM HGU through a neural mechanism that would increase GK transcription and translation in the morning. Second, it is well supported that insulin acts as a potent inhibitor of lipolysis, which in turn results in the improvement of whole-body glucose metabolism (44–49). We previously found that whereas selective AM hyperinsulinemia (accompanied by euglycemia) nearly completely suppressed circulating NEFA concentrations and enhanced PM HGU, selective AM hyperglycemia (with basal insulin) only reduced NEFA by 44% and did not enhance HGU (16). Thus, decreased NEFA could be an indirect mediator of the priming effect. Third, insulin inhibits glucagon secretion by the pancreatic alpha cell, which promotes glucose uptake by the liver (50–52). The present study, however, eliminates each of these indirect possibilities. Since arterial insulin levels increased similarly in both groups, brain insulin exposure was the same and lipolysis was equally suppressed. Likewise, differences in glucagon were prevented by somatostatin infusion with identical basal replacement of glucagon. Therefore, engaging insulin’s indirect effects in the morning was insufficient to bring about the priming of the liver for substantially enhanced PM HGU. Instead, a sufficient rise in insulin directly at the liver was key to improving glucose metabolism later in the day.

Skeletal muscle is the major site of insulin-dependent glucose uptake in the body under euglycemic conditions, providing a large tissue reservoir that can take up large amounts of glucose (53; 54). Although hyperinsulinemic-euglycemic clamps are often considered the gold standard for studies investigating the effects of insulin on glucose metabolism, they do not represent what occurs in normal physiology (21–25). In line with previous findings (55), regardless of the route of insulin delivery in the AM, MGU was largely responsible for the disposal of glucose during the euglycemic morning period (85-92% of total GIR). On the other hand, under meal-simulated conditions (hyperinsulinemia, hyperglycemia, and a negative arterial to portal vein glucose gradient), we have shown that liver glucose disposal can be as important as muscle, but only when insulin is delivered into the hepatic portal vein (19; 56). The current study extends this finding, demonstrating that portal vein insulin priming in the morning promotes the appropriate distribution of glucose disposal between the liver and muscle.

We also sought to understand the molecular changes that accompanied the profound enhancement of PM HGU and hepatic glycogen storage associated with AM insulin delivery. It is well known that GK acts as a gatekeeper to glucose entry into the liver (57; 58). Its transcription is rapidly and potently stimulated by insulin, remains elevated as long as insulin is present, and has been observed to reach maximal mRNA levels 4-8h after being stimulated with insulin (59–62). The half-life of glucokinase transcript and protein levels is not well-established in humans or dogs, although studies in rats suggest that GK mRNA decays rapidly with a half-life of 40-45 minutes (63; 64). With portal vein insulin delivery in the morning, there was a 3.5-fold elevation in GK mRNA at the beginning of the PM clamp which translated into a 2.7-fold increase in GK protein, effects that did not occur with AM peripheral insulin infusion. Thus, this augmentation in GK mRNA and protein in the AM Po Ins group prior to the onset of the PM clamp was likely due to an increase in hepatic insulin signaling throughout the AM clamp period, which is not captured in the hepatic tissue analysis 1.5h after the AM clamp ended. It should be noted that accurately measuring GK translocation from the nucleus to the cytoplasm, which is an important aspect of GK-mediated glucose uptake, is not feasible using our experimental model and methods (as previously described) (27). GS likewise plays a key insulin-mediated role in promoting liver glycogen storage (58). Compared to peripheral insulin delivery, infusion of insulin into the portal vein led to increased GS dephosphorylation and enzyme activity. GK expression and GS activity are reported to respond in a coordinated fashion to an insulin stimulus, as GK expression is a major site of control for the rate of glycogen synthetic flux (57; 65; 66). Thus, these data suggest that GK and GS were key players in the priming event caused by direct insulin action in the morning.

The present study demonstrates that morning portal vein insulin delivery enhanced the liver’s PM metabolic response compared to morning peripheral vein delivery. However, in the latter, hepatic insulin levels still increased 2-fold in the morning compared to basal insulin levels. Therefore, we also sought to determine whether peripheral insulin delivery was able to elicit any priming effect. To do so, for reference we compared the AM Pe Ins group to a previous study that had the same PM hyperinsulinemic-hyperglycemic clamp but mimicked “breakfast skipping” in the AM (morning insulin levels were basal) (39). While not statistically significant, mean PM HGU was 2.8 ± 0.3 mg/kg/min vs. 3.5 ± 0.3 mg/kg/min in the “breakfast skipping” vs. AM Pe Ins group, respectively. The effect of morning hyperinsulinemia on PM glucose metabolism was profoundly diminished when morning insulin was administered via a peripheral route, yet there was a tendency for an enhancement in afternoon HGU that was proportional to the insulin levels at the liver. The necessary hepatic insulin exposure required to prime the liver could be attained by increasing the morning rate of peripheral insulin delivery. However, this approach comes at the expense of increasing arterial hyperinsulinemia, which significantly increases the risk of hypoglycemia, and causes other metabolic defects (19; 21; 23; 24; 56; 67; 68).

There are few studies assessing the second meal phenomenon in the context of type 1 diabetes (69; 70). While it has been shown that subcutaneous insulin therapy can decrease the plasma glucose excursion following a second identical mixed meal, these individuals have a profound defect in net hepatic glycogen synthesis when compared to healthy individuals (69; 70). This may, in part, be associated with a loss of enhanced liver glucose metabolism caused by reduced direct hepatic insulin action. Thus, liver-targeted insulin therapy could help restore normal postprandial liver glucose uptake and storage in individuals with diabetes. Current approaches under investigation include intraportal islet transplantation, adjunct therapy, and oral and hepatopreferential insulin analogs (19; 71-74). In future studies, we will explore these therapeutic avenues to determine how targeting insulin to the liver in the morning might impact PM hepatic glucose metabolism during similar PM clamp conditions as outlined in this study.

In conclusion, our study highlights the critical importance of achieving appropriate hepatic insulin exposure in the morning to effectively prime the liver for increased HGU and hepatic glycogen synthesis later in the day. Morning peripheral insulin failed to induce this priming effect (at the rate used in this experiment), emphasizing the need to consider the route of insulin administration in therapeutic strategies. Future research should further explore the underlying mechanisms and clinical implications of these findings, particularly for optimizing insulin therapy in diabetes management.

## Supporting information

https://doi.org/10.6084/m9.figshare.26800804.v3

## Author Contributions

H.W., M.C.M., and A.D.C. participated in the design of experiments; H.W. and M.C.M. directed all experiments, collected, and interpreted data; B.F., K.Y., and M.S. participated in the experiments; B.F. and K.Y. were responsible for surgical preparation and oversight of animal care; H.W. and M.S.S. carried out biochemical and tissue analysis; H.W., D.S.E., and A.D.C. interpreted experimental results; H.W. prepared figures and drafted the manuscript; H.W., D.S.E., and A.D.C. edited and revised the manuscript; All authors approved the final version of the manuscript.

## Guarantor Statement

A.D.C. is the guarantor of this work and, as such, has full access to all the data in the study and takes responsibility for the integrity of the data and the accuracy of the data analysis.

## Conflict of Interest

A.D.C. has research contracts with Fractyl, Abvance, Fluidics, and Novo Nordisk, as well as grants from the NIH and the Helmsley Charitable Trust. A.D.C. is a consultant to Novo Nordisk, Paratus, Portal Insulin, Fractyl, and Thetis. No other individuals have conflicts of interest relevant to this article.

## Funding

This work was supported by National Institutes of Health (NIH) Grant R01DK131082. H.W. was supported in part by the Vanderbilt University Training Program in Molecular Endocrinology NIH grant 5T32DK007563. Hormone analysis was completed by Vanderbilt’s Hormone Assay and Analytical Services Core, supported by NIH grants DK020593 and DK059637. Vanderbilt’s Large Animal Core provided surgical expertise, supported by the NIH Grant DK020593.

## Prior Presentation

This work was presented at the American Physiology Summit 2024 meeting. The abstract can be viewed at doi.org/10.1152/physiol.2024.39.S1.470.

## Notes

https://doi.org/10.6084/m9.figshare.26800804.v3

