## Supplementary material for "Morning Engagement of Hepatic Insulin Receptors Improves Afternoon Hepatic Glucose Disposal and Storage": https://doi.org/10.6084/m9.figshare.26800804.v3


### Western Blotting

The following primary antibodies were used in western blotting analysis: pAkt (#9271, Cell Signaling Technology, Danvers, MA), tAkt (#9272, Cell Signaling Technology), pGS (#98348, Cell Signaling Technology), tGS (#3893, Cell Signaling Technology), pGSK3β (#9336, Cell Signaling Technology), tGSK3β (#9315, Cell Signaling Technology), and GK (#sc-17819, Santa Cruz Biotechnology). Anti-mouse and Anti-rabbit IgG (H+L), HRP conjugate secondary antibodies were purchased from Promega (Madison, WI).

Protein samples were loaded in order by group for all western blot images. The manuscript expresses these data as phosphorylated proteins relative to total protein for Akt, GS, and GP. GK is expressed over Cyclophilin B. The image for each respective protein can be viewed below.

### AM Po Ins (No PM Clamp and PM Clamp) Western Blot Images

Phosphorylated Akt (pAkt)

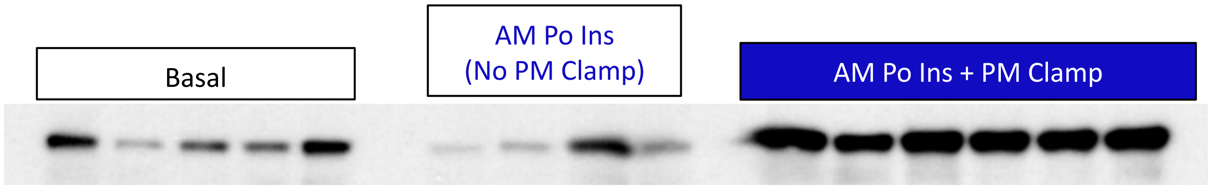


Total Akt (tAkt)

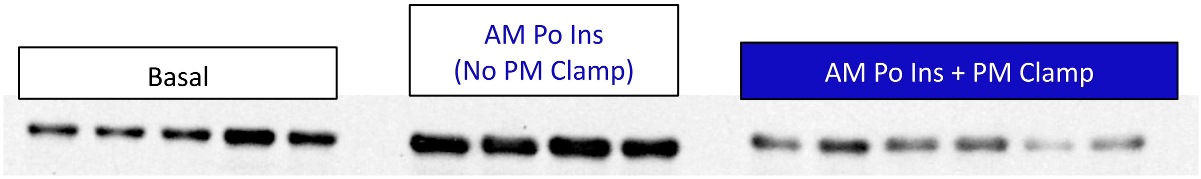


GK

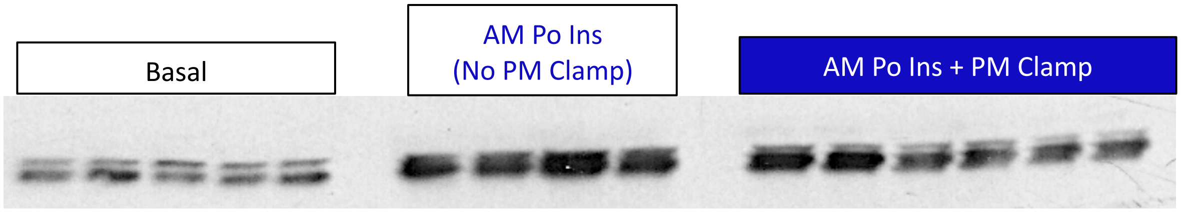


Cyclophilin B

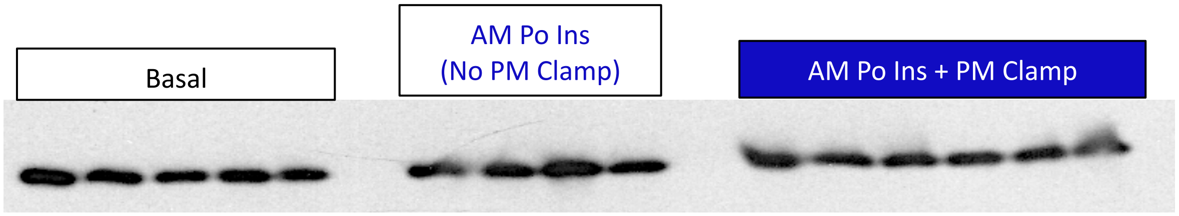


Phosphorylated GS (pGS)

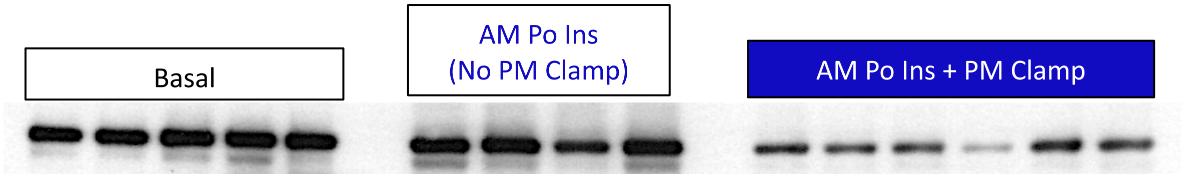

Total GS (tGS)

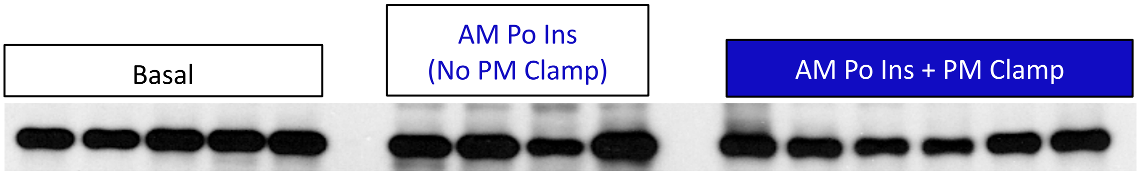


Phosphorylated GP (pGP)

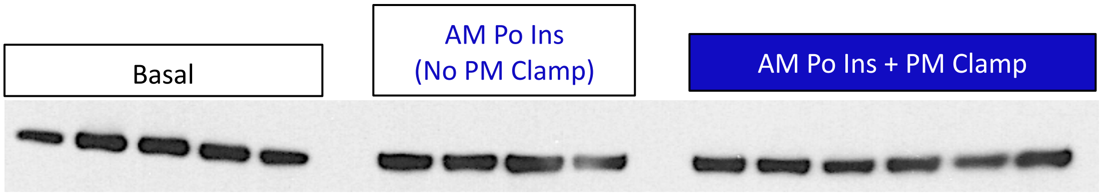


Total GP (tGP)

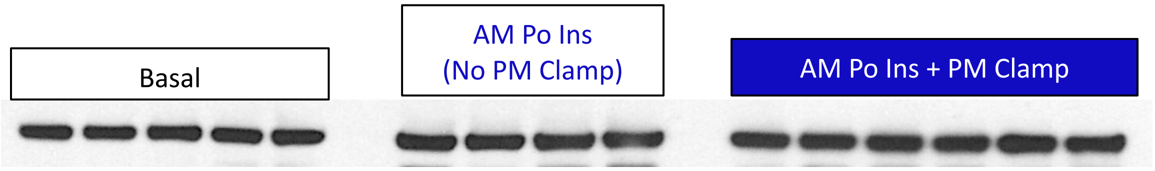


**AM Pe Ins (No PM Clamp and PM Clamp) Western Blot Images**

Phosphorylated Akt (pAkt)

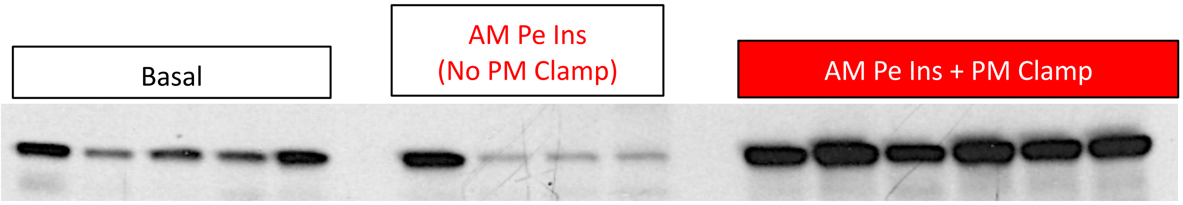

Total Akt (tAkt)

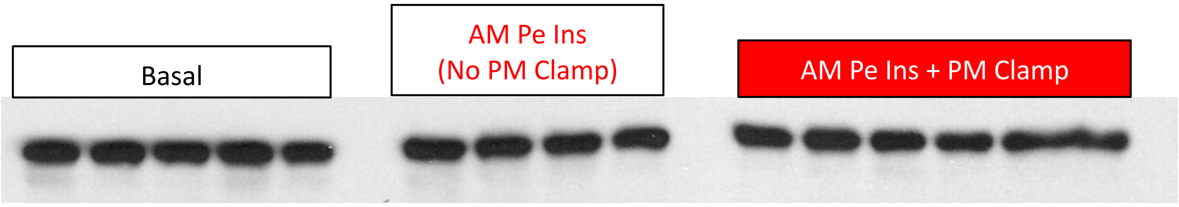


GK

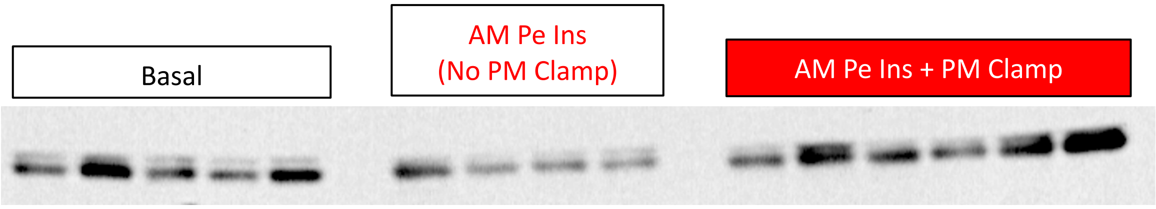


Cyclophilin B

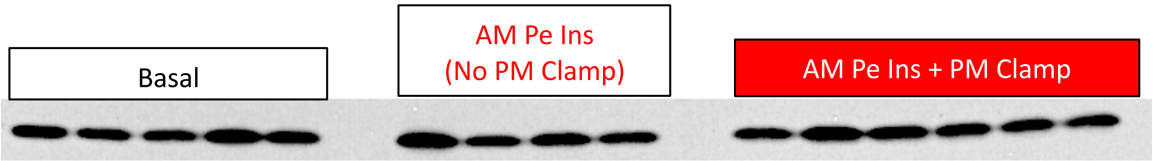


Phosphorylated GS (pGS)

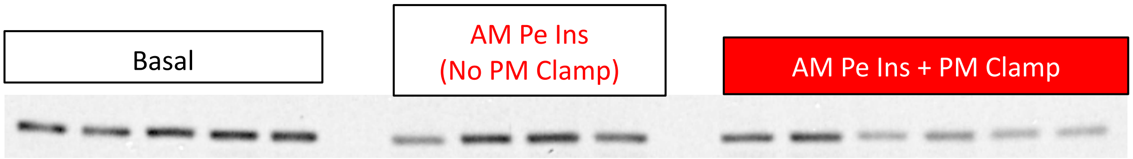


Total GS (tGS)

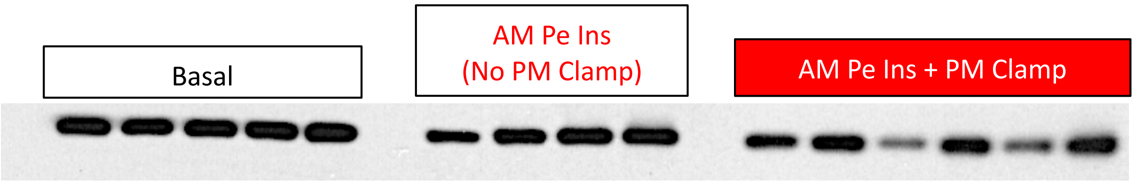

Phosphorylated GP (pGP)

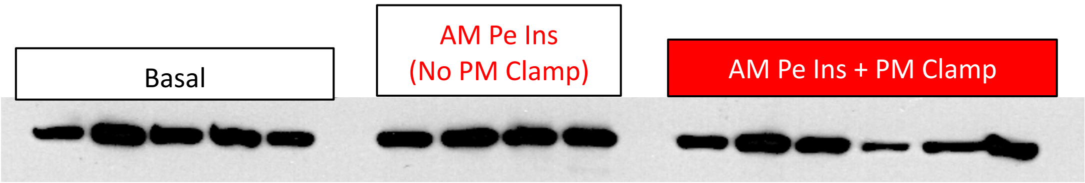


Total GP (tGP)

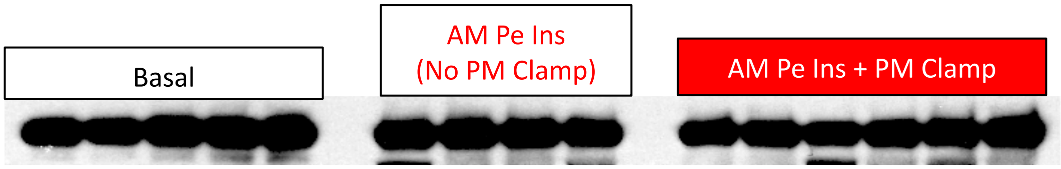
